# Numerical evaluation reveals the effect of branching morphology on vessel transport properties during angiogenesis

**DOI:** 10.1101/2020.10.13.337295

**Authors:** Fatemeh Mirzapour-shafiyi, Yukinori Kametani, Takao Hikita, Yosuke Hasegawa, Masanori Nakayama

## Abstract

Blood flow governs transport of oxygen and nutrients into tissues. Hypoxic tissues secrete VEGFs to promote angiogenesis during development and in tissue homeostasis. In contrast, tumors enhance pathologic angiogenesis during growth and metastasis, suggesting suppression of tumor angiogenesis could limit tumor growth. In line with these observations, various factors have been identified to control vessel formation in the last decades. However, their impact on the vascular transport properties of oxygen remain elusive. Here, we take a computational approach to examine the effects of vascular branching on blood flow in the growing vasculature. First of all, we reconstruct the 3D vascular model from the 2D confocal images of the growing vasculature at P6 mouse retina, then simulate blood flow in the vasculature, which is applied for the gene targeting mouse models causing hypo- or hyper-branching vascular formation. Interestingly, hyper-branching morphology attenuates effective blood flow at the angiogenic front and promotes tissue hypoxia. In contrast, vascular hypo-branching enhances blood supply at the angiogenic front of the growing vasculature. Oxygen supply by newly formed blood vessels improves local hypoxia and decreases VEGF expression at the angiogenic front during angiogenesis. Consistent with the simulation results indicating improved blood flow in the hypo-branching vasculature, VEGF expression around the angiogenic front is reduced in those mouse retinas. Conversely, VEGF expression was enhanced in the hyper-branching vasculature in the mouse retina. Our results indicate the importance of detailed flow analysis in evaluating the vascular transport properties of branching morphology of the blood vessels.

**Author Summary:** Blood vessels are important for the transport of various substances, such as oxygen, nutrients, and cells, to the entire body. Control of blood vessel formation is thought to be important in health and disease. In the last decades, various factors which regulate blood vessel branching morphology have been identified. Gene modification of some of these identified factors results in hyper-branching of the vasculature while others cause hypo-branching of the vessel. Given the importance of the transport property of the blood vessel, it is important to examine the effect of these identified factors on the transport property of the affected vascular morphology. In line with these facts, we reconstruct 3D vessel structures from 2D confocal microscopy images. We then simulate blood flow in the structures numerically. Interestingly, our results suggest vessel network complexity negatively affects the blood perfusion efficiency and tissue oxygenation during angiogenesis. Thus, our results highlight the importance of flow analysis considering the detailed 3D branching pattern of the vascular network to quantitatively evaluate its transport properties.

## Introduction

Upon tissue hypoxia, proangiogenic factors such as vascular endothelial growth factors (VEGFs) are secreted to induce new blood vessel formation from existing vessels, termed angiogenesis [1]. While it is important to promote angiogenic vessel growth for tissue homeostasis, the vessel formation in tumors can give transformed cells better access to nutrients and oxygen [2–4]. Transformed cells hijack blood vessels for metastasis to a distant tissue [5]. In the last decades, various studies have identified critical factors controlling endothelial proliferation and vascular branching [6–10]. In line with these observations, anti-angiogenic therapy aimed to starve tumor cells of nutrients and oxygen by reducing tumor vascularization. However, the outcome of the treatment was more limited than expected [11, 12]. Reducing blood vessel formation in tumor tissues is thought to enhance ischemia to induce tumor resistance against chemotherapies as well as to restrict drug delivery [6, 13]. However, the effects of altered vascular branching on blood flow distribution and the oxygen transport property have not been investigated yet.

Here, we introduced numerical evaluation of branching morphology of mouse retinal growing vasculatures on vessel transport properties. Interestingly, hyper-branching morphology attenuates effective blood flow at the angiogenic front *in silico* and fails to improve tissue hypoxia *in vivo*. In contrast, hypo-branching morphology enhances blood supply at the growing vasculature. Consistently, VEGF expression of the angiogenic front region was efficiently improved, suggesting better oxygen supply at the region. Our results indicate the importance of evaluating branching networks of the vasculature by transport property of the blood vessel.

## Results

### Image acquisition and processing of vascular network

First, we reanalyzed the postnatal day 5 (P5) vasculature of the endothelial cell (EC) specific tamoxifen inducible gene deficient mice. As the models for the hypo- and hyper-branched vasculature, we employed *lox* P-flanked *Prkci* [14] and *lox* P-flanked *Foxo1* [15] mice crossed with *pdgfb*-icre mouse. Endothelial proliferation is modulated via controlling c-Myc expression in both model mice [15, 16]. After tamoxifen injection from P1 to P3 to induced effective gene deletion, mouse retina was harvested and stained with an anti-ICAM-II antibody the marker of the inner lumen of the blood vessels. Stained vasculatures were visualized by confocal microscopy. To measure the property of the vasculature, morphometric measurements were introduced (Fig. 1A). Consistent with the previous report [16], the retinal vasculature in the *Foxo1* EC specific inducible knock out (*Foxo1*^iΔEC^) showed increased vascular density, vessel length density and branching index (Fig. 1B to E), while the same factors in the *Prkci* EC specific inducible knock out (*Prkci*^iΔEC^) were decreased (Fig. 1F to I).

**Figure 1.**
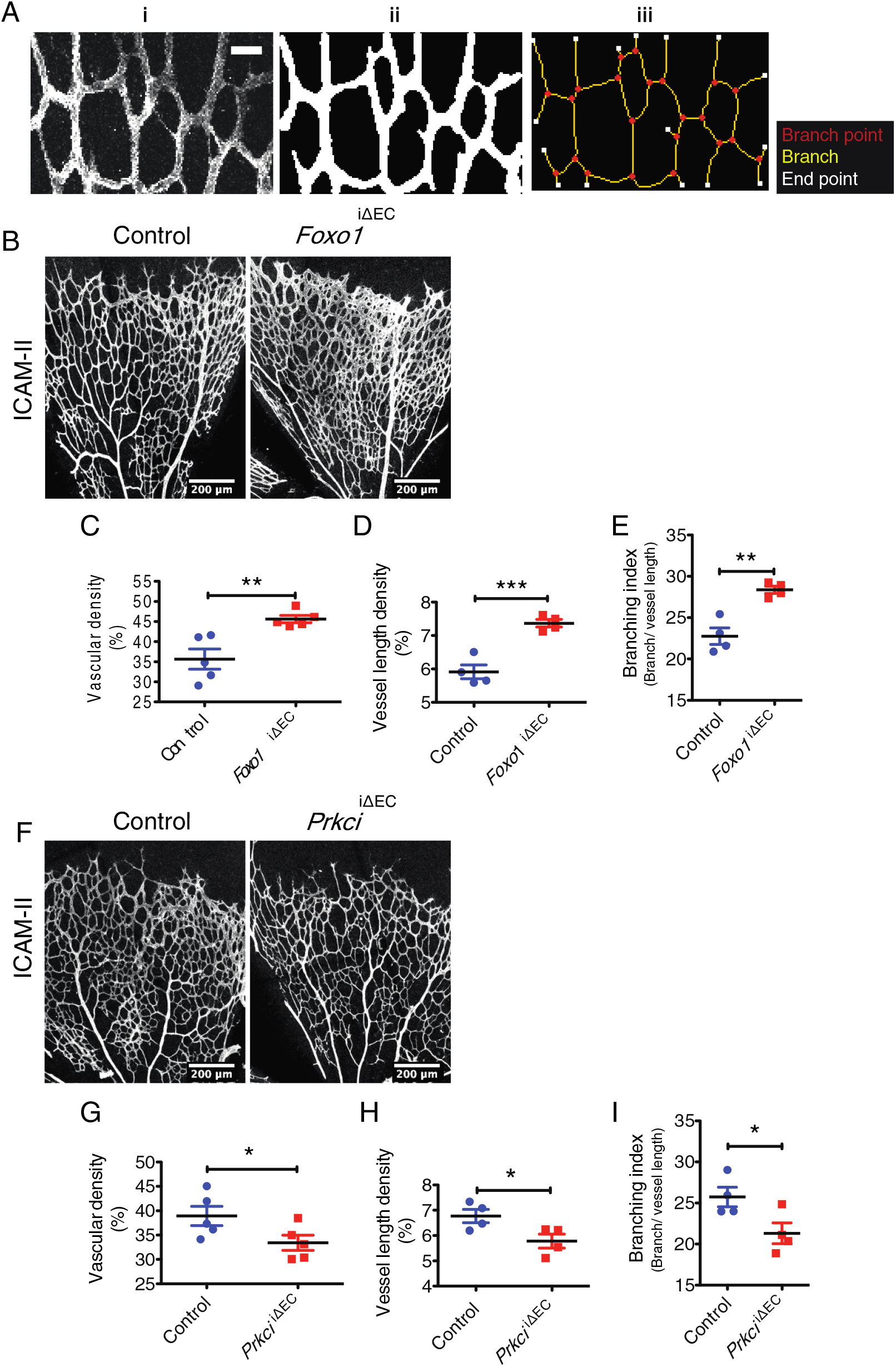
Characterization of the hyper- and hypo-branching vascular models. **A)** Grayscale image of a sample subset of the retinal microvasculature stained for ICAM-II (i), converted into the vessel mask (ii) and used for skeleton analysis (iii); Scale bar represents 30 μm. **B)** Staining of ICAM-II in control and *Foxo1*^*i*ΔEC^ mouse retinae at postnatal day 5 (P5). **C)** Quantification of vascular density (percentage of area occupied by vessels), **D)** vessel length density (percentage of area occupied by skeletonized vessel) and **E)** branching index (branch point/ mm of vessel length) in control and *Foxo1*^*i*ΔEC^ retina at P5. **F)** Staining of ICAM-II in control and *Prkci*^iΔEC^ mouse retinae at P5. **G)** Quantification of vascular density (percentage of area occupied by vessels), **H)** vessel length density (percentage of area occupied by skeletonized vessel) and **I)** branching index (branch point/ mm of vessel

### 3D reconstruction of vascular network

Next, we reconstructed the three-dimensional (3D) vascular model from the two-dimensional (2D) confocal images. The obtained RGB images were converted to black and white binarized images (‘vessel mask’) using MATLAB Image Processing Toolbox (Fig. 2A), where white and black pixels corresponded to the regions of the blood vessel and the surrounding tissue, respectively. The binarized vessel structure was then projected onto a *x-y*|_z=0_ plane in the 2D Cartesian coordinate system (Fig. 2B). For all grid points within the blood vessel the shortest distance to the vessel wall was calculated in the 2D structure, and then a 3D sphere with the diameter obtained at each grid point was placed in a 3D Cartesian coordinate system as schematically shown in Fig. 2C. The envelope of all the spheres was then used to define the 3D blood vessel structure (Fig. 2D). After reconstructing the 3D vascular structure, a signed distance function, which is commonly referred to as a level-set function [17], was computed at every grid point in both vessel and tissue regions. The obtained level-set function was integrated to an in-house solver for the blood flow.

**Figure 2.**
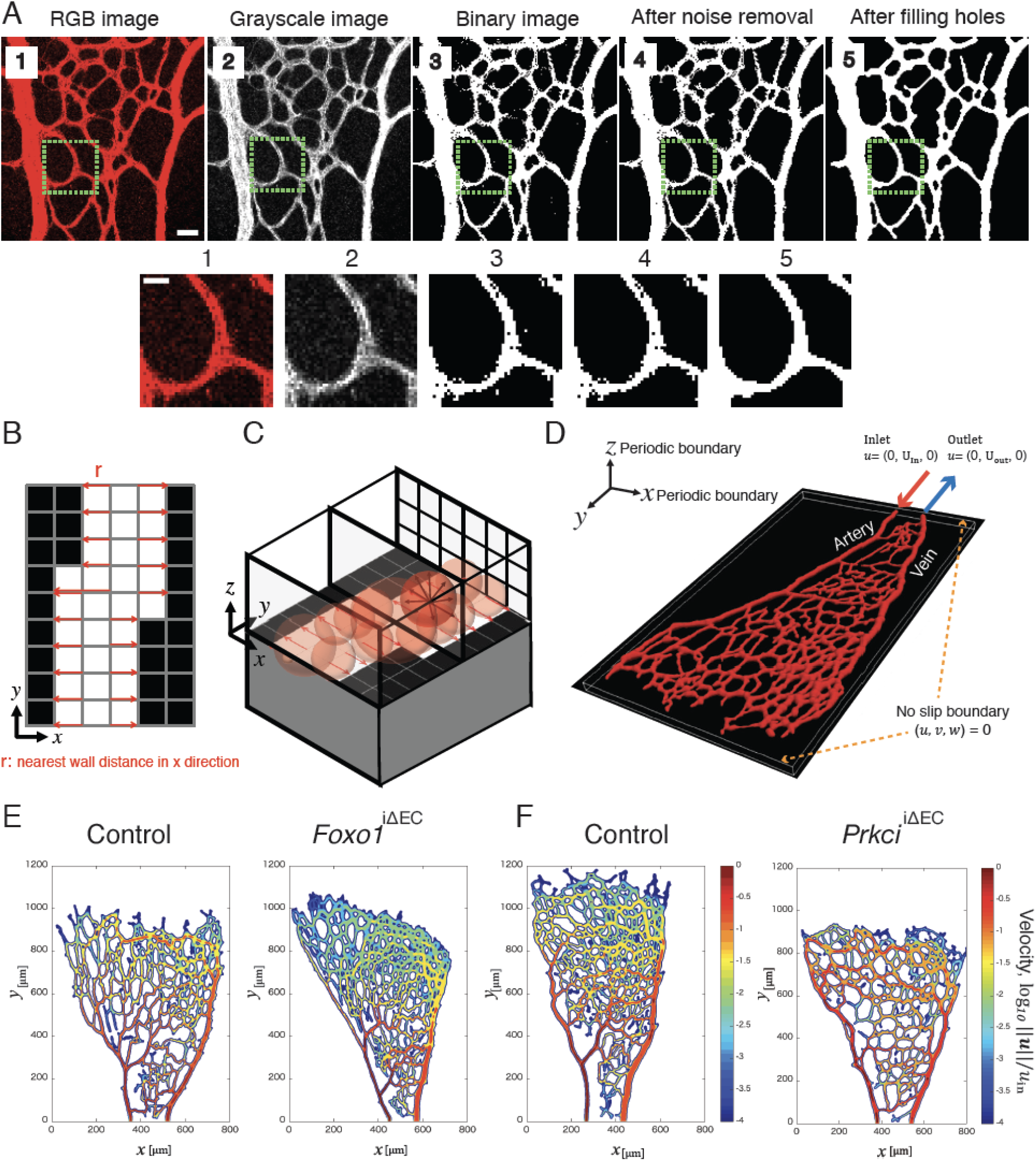
Image processing and blood flow simulation from the growing mouse retinal vasculature. **A)** Generation of binary vessel mask from RGB images. Higher magnification images of indicated areas in the upper panels are shown in the lower panels. Scale bar in the upper panel, 30 μm; 10 μm in the lower panel. **B)** Schematic of the searching process for the nearest solid wall (black pixel) on each pixel point and defining the local radius *r* (pixel distribution is projected on Cartesian grid coordinate 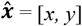). **C)** The local radius *r* for creating 3D vascular structure in a 3D Cartesian coordinate system, ***x*** = [*x, y, z*], extended from 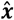. The region with ‖*x*‖ < *r* is defined as blood vessel, while the rest is a surrounding tissue. **D)** Sample reconstructed 3D model of the retinal vasculature with indication of the inlet and outlet in Cartesian coordinate system **E, F)** Visualization of the normalized velocity amplitude in logarithmic scale (‘log_10_‖*u*‖/*u*_*in*_’) on the central *x-y* plane (along the *z* axis) for *Foxo1*^*i*ΔEC^ (E) and *Prkci*^iΔEC^ mice (F) with control.

### Numerical simulation of blood flow

As a suspension of erythrocytes in plasma, blood becomes a shear-thinning fluid due to rouleaux formation [20]. Rouleaux, aggregation of red blood cells (RBC), causes increased blood viscosity due to increased effective volume of RBCs [20]. Thus, blood behaves as a non-Newtonian fluid as its viscosity decreases with applied shear. However, under a sufficiently high shear rate (> 150 s^−1^, [21]), blood flow can be accurately modeled as Newtonian fluid with a constant viscosity [22]. Nagaoka et al. measured averaged values of 632 to 1539 s^−1^ for shear rate in human retinal first arteriole and venule branches [23]. Moreover, Windberger et al. [24] measured species-specific effect of shear rate on blood viscosity and erythrocytes aggregation within low shear rates of 0.7, 2.4 and 94 s^−1^. They reported different viscosity values and RBC aggregation index among different mammalian species: horse, pig, dog, cat, rat, cattle, sheep, rabbit and mouse. In low shear rate regime (0.7 and 2.4 s^−1^), they found lower shear dependent viscosity enhancement in cattle, sheep, rabbit and mouse, as compared to horse, rat, pig, dog and cat. Compared to other species, erythrocyte aggregation (EA, measured using four different methods) in mouse was found low and, in some methods, undetectable. At high shear rate (94 s^−1^) they found the EA destroyed and the RBCs orientated to the flow direction. Based on these results, a simple Newtonian behavior is accounted to retinal flow in this model.

*In vivo* measurements have shown systolic and diastolic flow rate variations in human [25] and mouse [26] retinal vasculature. The influence of pulsation on flow is define by the Womersley number, first described in 1955 [27]. This dimensionless number can be considered as the time ratio between the pulsation period and the time-scale for the wall information propagates to the bulk fluid via fluid viscosity, and therefore it determines the pulsations effects on the velocity profile. Specifically, when the Womersley number (α) is sufficiently low (≤ 1), there is enough time for a velocity profile to develop during each cycle (viscous-dominated flow), so that the resultant flow can be considered as a quasi-steady flow. While, for large values of α (≥ 10), the transient interaction between pulsation and blood flow plays an essential role and thereby the unsteady flow analysis is crucial. The Womersley number is defined by the following formula:

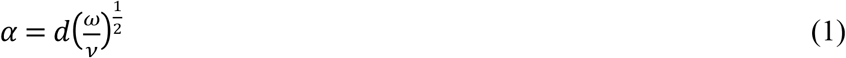

Where *d*, ω and ν are the vessel diameter, the angular frequency for a heart rate and the kinematic fluid viscosity, respectively. Using in vivo reported values for mouse as such: *d* ~ 10^−5^ [m], ω ~1 [s^−1^], and ν ~10^−6^ [m^2^/s] [28–30], in neonatal mouse retina, α becomes as low as ~10^−2^ (≪ 1). Hence, the flow in retina model can be considered in viscous-dominated regime and this validates the quasi-steady assumption mentioned above. The other blood rheological properties such as the Fåhræus–Lindqvist effects, and the elastic effects of a vessel wall are not taken into account in the present model. Consequently, the flow is assumed to be incompressible, Newtonian and steady. The governing equations for the blood flow are given by the following Navier-Stokes and continuity equations:

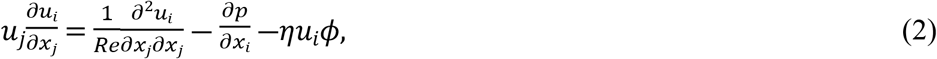

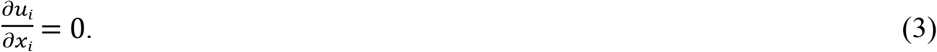

Here, *u*_*i*_, *p* and *Re* denote the velocity component in the *i*-th direction, the static pressure and the Reynolds number, respectively. The inlet bulk mean velocity 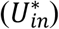 and the diameter of the inlet artery 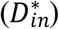 were used for non-dimensionalization in Eqs. (2, 3). Here, the superscript of * represents a dimensional value, whereas a quantity without a superscript indicates a dimensionless quantity. The Reynolds number, expressing the ratio of the inertial and viscous forces, was defines as: 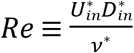. In this study, *Re* was set to be 0.1 based on 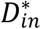 and the kinematic viscosity and a typical blood flow velocity in an artery reported in [24, 28].

The current code is based on a volume penalization method [18], which is categorized as immersed boundary techniques [19]. The main advantage of the present approach is that an arbitrary 3D structure can be embedded in 3D Cartesian computational grids, so that there is no need to generate boundary-fitted grids for each geometry. Meanwhile, it has relatively slow convergence due to smearing of the fluid-solid boundary, so that we performed grid convergence studies to confirm that the present conclusions are not affected by further grid refinement (Fig. S1). The last term on the right-hand-side of Eq. (2) corresponds to an artificial body force term introduced in the volume penalization method (VPM) in order to realize a no-slip condition at a fluid-solid boundary [15]. Computational grid points were uniformly distributed in space and the spatial grid resolutions were set to be 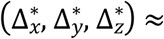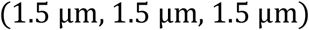. The dimensions of the computational domain for each model are listed in Table1. The dimensions of the computational domain depend on each sample, while the computational resolutions were kept constant for all the cases.

**Table 1.** Domain size and grid resolution for every model structure.

| Name | Domain size<br>$L_x^* \times L_y^* \times L_z^*$ [μm] | $D_{in}^*$ [μm] | Grid size<br>[μm] | Number of grid points<br>$N_x \times N_y \times N_z$ |
| --- | --- | --- | --- | --- |
| *Foxo1* CTRL-1 | 800.30 × 1072.10 ×<br>33.22 | 12.08 | 1.51 | 530 × 710 × 22 |
| *Foxo1*<sup>iΔEC</sup>-1 | 755.00 × 1102.30 ×<br>36.24 | 12.08 | 1.51 | 500 × 730 × 24 |
| *Foxo1* CTRL-2 | 588.90 × 936.20 × 36.24 | 12.08 | 1.51 | 390 × 620 × 24 |
| *Foxo1*<sup>iΔEC</sup>-2 | 619.10 × 694.60 × 36.24 | 10.57 | 1.51 | 410 × 460 × 24 |
| *Foxo1* CTRL-3 | 694.60 × 1087.20 ×<br>30.20 | 13.59 | 1.51 | 460 × 720 × 20 |
| *Foxo1*<sup>iΔEC</sup>-3 | 936.20 × 1011.70 ×<br>33.22 | 13.59 | 1.51 | 620 × 670 × 22 |
| *Prkci* CTRL-1 | 664.40 × 1208.00 ×<br>36.24 | 13.59 | 1.51 | 440 × 800 × 24 |
| *Prkci*<sup>iΔEC</sup>-1 | 830.50 × 966.40 × 33.22 | 15.10 | 1.51 | 550 × 640 × 22 |
| *Prkci* CTRL-2 | 845.60 × 1253.30 ×<br>36.24 | 13.59 | 1.51 | 560 × 830 × 24 |
| *Prkci*<sup>iΔEC</sup>-2 | 543.60 × 1208.00 ×<br>30.20 | 10.57 | 1.51 | 360 × 800 × 20 |
| *Prkci* CTRL-3 | 588.90 × 951.30 × 33.22 | 12.08 | 1.51 | 390 × 630 × 22 |
| *Prkci*<sup>iΔEC</sup>-3 | 588.90 × 875.80 × 27.18 | 10.57 | 1.51 | 390 × 580 × 18 |
| *Cases with finer resolution for verification of grid resolutions* |  |  |  |  |
| *Foxo1* CTRL-1F | 800.30 × 1072.10 ×<br>33.22 | 12.08 | 0.76 | 1060 × 1420 × 44 |
| *Foxo1*<sup>iΔEC-1F</sup> | 755.00 × 1102.30 ×<br>36.24 | 12.08 | 0.76 | 1000 × 1460 × 48 |
| *Prkci*<sup>CTRL-1F</sup> | 664.40 × 1208.00 ×<br>36.24 | 13.59 | 0.76 | 880 × 1600 × 48 |
| *Prkci*<sup>iΔEC-1F</sup> | 830.50 × 966.40 × 33.22 | 15.10 | 0.76 | 1100 × 1280 × 44 |

The computational domains are schematically shown in Figure 2A. Uniform velocity profiles were applied at the inlet and outlet boundaries, *U*_*in*_ and *U*_*out*_, respectively. The velocity at the outlet was determined such that the fluid volume is strictly conserved throughout the structure. At the outer boundaries of the computational domain, a Neumann boundary condition was imposed for the pressure.

### Simulation results

The obtained velocity fields showed distinct flow distributions for hyper- and hypo-branched structures (Fig. 2E and F). In general, the flow distribution was un-even in the *Foxo1*^iΔEC^ vessel network comparing to that in control. Specifically, the velocity around the peripheral edge (angiogenic front) is drastically attenuated due to the hyper-branching. Conversely, the hypo-branched *Prkci*^iΔEC^ network results in a more evenly distributed flow throughout the entire network including the peripheral edge. Next, to quantify the amount of flow transported to the peripheral regions, we introduced a cylindrical coordinate system (*r*-*θ*) with its origin at the first branching point from either the artery or the vein near the inlet (Fig. 3A). Here, *r* and *θ* represent the distance from the origin and the azimuthal direction, respectively. Accordingly, the local flow velocity vector was decomposed in to the radial and azimuthal components, i.e., *u*_*r*_ and *u*_*θ*_ (Fig. 3A). The left figure of Figure 3B shows the spatial distribution of the absolute local velocity ‖*u*‖, normalized by the inlet velocity *u*_in_, while that of the normalized azimuthal flow velocity, *u*_*θ*_/*u*_in_, is depicted in the right figure of Figure 3B. To evaluate the perfusion efficiency of each structure, the entire vascular network was divided into two regions, i.e., angiogenic front and inner plexus (Fig. 3C and D). The angiogenic front was defined as the region of 0.7*R*_*max*_ < *r* <*R*_*max*_, where *R*_*max*_ is the distance of the farthest blood vessel from the origin. The current definition of the angiogenic front is based on the previous observation that c-Myc expression and the proliferation of ECs are active in the region [15, 16]. Here, *R*_*max*_ is the distance of the farthest blood vessel from the origin. The azimuthal flow rates (*Q*_*θ*_) for the two regions were separately calculated through volume integration of *u*_*θ*_. This way, the portion of the blood flow transported to the angiogenic front can be quantitatively evaluated. While the vessel density was increased in the *Foxo1*^iΔEC^ mice, *Q*_*θ*_ at the angiogenic front was significantly decreased in *Foxo1*^iΔEC^ mice (Fig. 3E and G). In contrast, it was increased for *Prkci*^iΔEC^ mice compared to the control littermates, although vascular branching was decreased (Fig. 3F and H).

**Figure 3.**
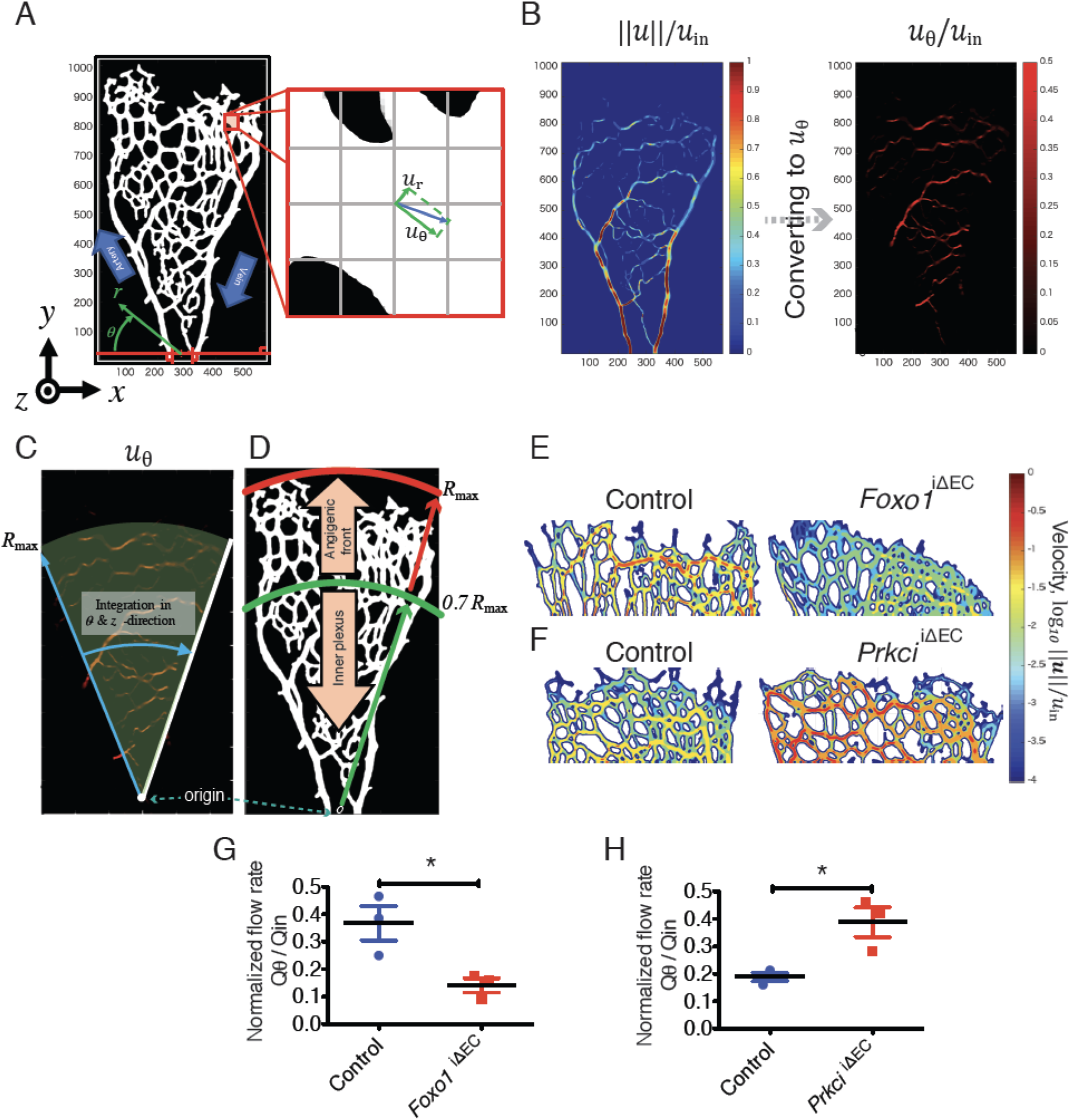
Quantification of blood flow at the angiogenic front. **A)** Coordinate transformation from the Cartesian system (*x, y*) to the cylindrical system (*r, θ*). **B**) The velocity field transformed from the Cartesian coordinate system to the cylindrical one. **C**) Distribution of the azimuthal velocity *u*_*θ*_ between the artery and the vein. **d**) Schematic of defining the angiogenic front. The entire vasculature is separated into two regions: the inner plexus and the angiogenic front **E)** The velocity amplitude distribution at the angiogenic front in control and *Foxo1*^*i*ΔEC^ structures at P5, and **F)** in control and *Prkci*^iΔEC^ structures at P5. **G, H)** The normalized azimuthal flow rate at the angiogenic front of the *Foxo1*^*i*ΔEC^ (G) and *Prkci*^iΔEC^ mice (H) with control. Data represent mean ± S.E.M. two-tailed unpaired t-test *p < 0.05 (n ≥ 3).

### Experimental Validation

The distinct blood flow distributions for *Foxo1*^iΔEC^ and *Prkci*^iΔEC^ mice should have significant impacts on their transport properties. To validate our numerical results, we observed VEGF expression with an anti-VEGF antibody in the mutant retinae, which reflects the hypoxic status of the tissue. During angiogenesis, VEGF is expressed around the angiogenic front. Once the tissue is vascularized, local hypoxia is improved by oxygen supply with the newly formed blood vessels to downregulate VEGF expression [1]. Consistent with the decreased flow rate at the angiogenic front in the *Foxo1*^iΔEC^ mice, VEGF expression around the angiogenic front were significantly increased compared to that of the control (Fig. 4A and C). Conversely, VEGF expression in *Prkci*^iΔEC^ mice around the angiogenic front region was decreased compared to that of the control (Fig. 4B and D).

**Figure 4.**
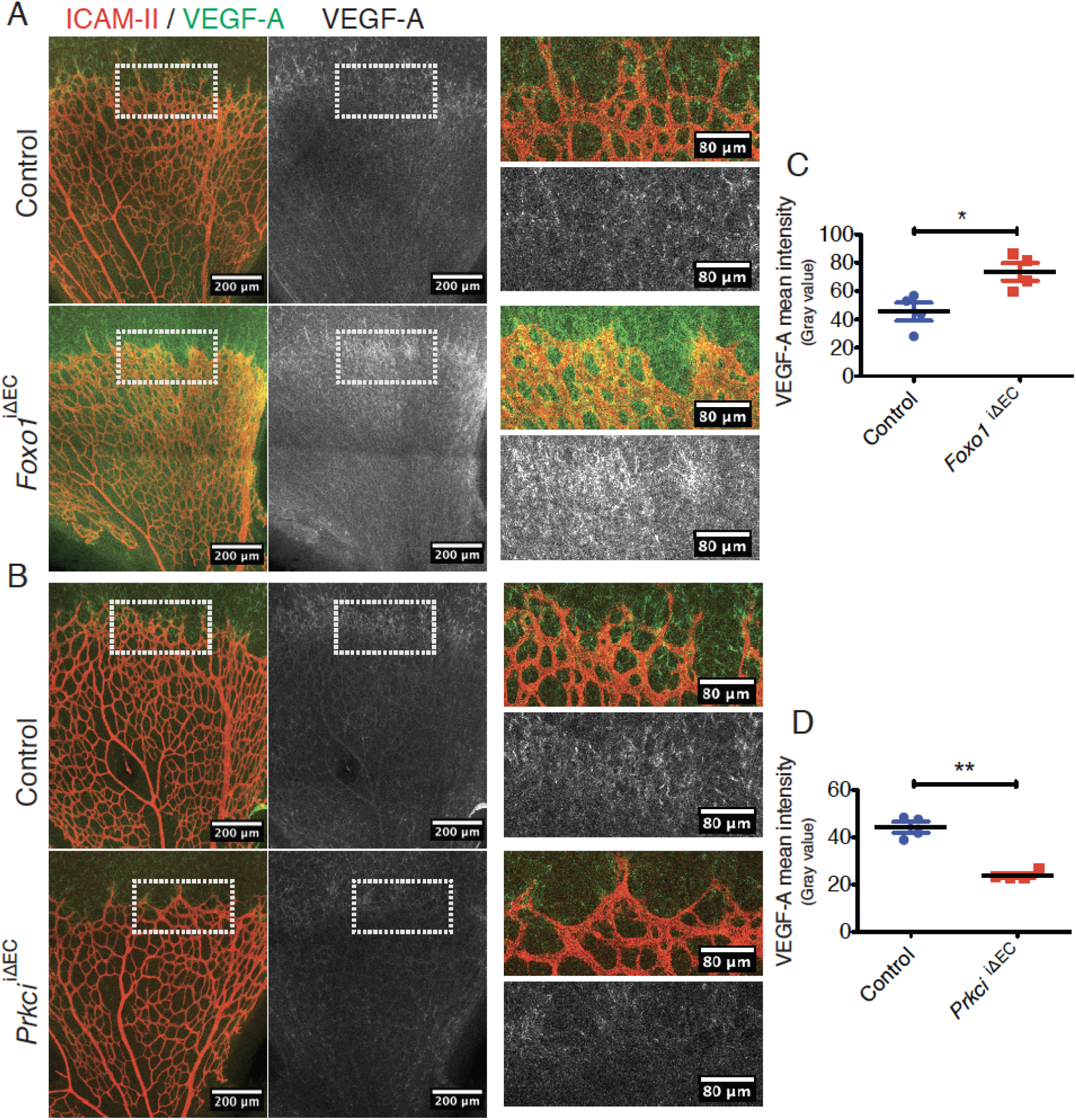
The effect of the hyper- or hypo-branching vasculature on VEGF expression during angiogenesis. **A, B)** Staining of ICAM-II (red) and VEGF-A (green and gray) in the *Foxo1*^*i*ΔEC^ (A) and *Prkci*^iΔEC^ (B) mouse retina with control. Higher magnification images of the indicated areas in the left panel are shown in the right panel. **C, D)** Quantification of VEGF-A signal intensity at angiogenic front in (A) and (B) respectively. Data represent mean ± S.E.M. two-tailed unpaired t-test *p < 0.05 **p < 0.01 (n ≥ 4).

## Discussion

While suppression of blood vessel formation has been expected to have a negative impact on blood supply, our results indicate the opposite trend. Namely, the suppression of vessel branching enhances blood flow, and thereby oxygen transport at the angiogenic front. Conversely, the enhancement of blood vessel formation attenuates blood flow at the angiogenic front. The present results underline the importance of detailed flow analysis considering a complex microvascular structure for evaluating its transport properties. In clinical applications, it has been reported that combining anti-angiogenic drug and chemotherapy statistically improves the progression-free survival rate [31]. This suggests that the normalization of the vascular network around tumors could contribute to delivering the medicine to the tumors by enhancing blood supply. Recently, the nanoparticulate drug delivery system has attracted much attention due to its potential to further improve the therapeutic efficacy. Since it is known that the transport of nanoparticles in vasculature is significantly affected by their sizes and shapes [32], the detailed analysis of blood flow and associated mass transfer becomes ever more needed. Computational fluid dynamics of blood flow in vascular networks should be a key for optimizing the shape, size and chemical properties of nanoparticles in drug delivery and even for the designing and control of microrobots for future medical applications [33].

Although flows in large vessels such as a coronary artery can be essentially modelled as homogeneous Newtonian fluid, flows in the microcirculation are strongly affected by the complex interactions among plasma flow, interstitial flow, complex geometry of branching patterns and the dynamics of blood cells whose size is close to the blood vessel diameter [34]. In order to accurately reproduce such multi-scale and multi-physics phenomena, 3D flow simulation is necessary. Due to its large computational cost, however, most existing studies relies on simplified 1D analysis [35, 36], whereas studies applying 3D analysis for vascular network are still limited [37]. In the present study, a new approach to implement a complex structure of vascular network into 3D flow simulation was introduced. By representing an arbitrary complex 3D network structure with the level-set function, the structure was immersed in 3D cartesian coordinate by the volume penalization technique. This has two major advantages: First, the grid generation which is required in flow simulation with a body-fitted coordinate system can be omitted, so that it becomes quite straightforward to simulate flows in different geometries. Second, since the present scheme deploys structured grids in both vessel and tissue regions, flow coupling between the blood and interstitial flows and mass transport between blood and tissue can be solved in a unified manner. The latter is particularly important when the transport of oxygen or nanoparticles from blood flow to surrounding tissues will be considered in future work.

## Materials and Methods

### Mice breeding

All animal experiments were conducted according to the protocols approved by the local animal ethics committees and authorities (Regierungspräsidium Darmstadt) and institutional regulations. As previously described, transgenic *Pdgfb*-iCre mice were bred into lines of animals containing a LoxP-flanked *Prkci* [14] and LoxP-flanked *Foxo1* [15]. Intraperitoneal injections of tamoxifen (sigma, T5648) from postnatal day1 (P1) to P3 was used to induce activation of Cre in neonatal mice. The phenotype of the mutant mice was analyzed at P5 and tamoxifen injected Cre negative littermates were used as controls.

### Retina staining

For retina staining, eyeballs were fixed for 20min in 2% Paraformaldehyde (PFA; Sigma, P6148) at room temperature (RT). Afterwards, retinas were dissected in PBS and fixed for 30min in 4% PFA on ice. Next, they were washed three times with PBS and permeabilized and blocked for 2hr at RT in blocking buffer (BB): 1% fetal bovine serum (FBS; Biochrom GmbH, Berlin, Germany), 3% Bovine Serum Albumin (BSA; Sigma, A2153), 0.5% Triton™X-100 (Sigma, T8787), 0.01% Na deoxycholate (Sigma, D6750) and 0.02% Na Azide (Sigma, S8032) in PBS on rocking platform. Then they were incubated with primary antibodies (anti-ICAM-II (BD Pharmingen™, 553326, 1:100), anti- Collagen-Type IV (Collagen-IV) (Bio-RAD, 2150-1470, 1:400) and anti-VEGF164 (R&D Systems, AF-493-NA, 1:100) in 1:1 BB/PBS), overnight at 4°C on rocking platform. Retinas were then washed four times for 30 min in PBS/ 0.2% Triton™X-100 (PBT) at RT and incubated with Alexa Fluor conjugated secondary antibodies (Invitrogen, 1:500) in 1:1 BB/PBS for 2hr at RT. After another four times of washing with PBT, retinas were radially cut into four lobes and flat-mounted onto slides using Fluoromount-G mounting medium (Southern Biotech, 0100-01).

### Image processing and statistical analysis

In experiments using KO animal models, data are derived from three independent experiments (three sets of mutant mice and control littermates). The data are presented as mean ± S.E.M. All statistical analyses were carried out using Prism software (GraphPad, CA); p<0.05 was considered as significantly different. Quantification of VEGF-A signal intensity was carried out by measuring mean gray value using ImageJ (1.52o). Morphometric analyses of the retinal vasculature were assessed using Fiji (2.0.0-rc-69/ 1.52n) Vessel Analysis plugin, Fiji Skeletonize plugin and Fiji Skeleton Analyzer. Volocity (Perkin Elmer, MA), Photoshop CS, Illustrator CS (Adobe), ImageJ and Fiji software were used for image processing in compliance with general guidelines for image processing.

### Numerical scheme

#### Spatial discretization

An energy conservative second-order central finite difference scheme (CDS) was applied for the convection term on the left-hand-side of Equation 2 on the staggered grid [38]. The diffusion term on the right side of Equation 2 was also discretized by a conventional second-order central finite difference scheme.

#### Temporal marching

A Simplified Marker and Cell, SMAC, [39] method was implemented to decouple the pressure in the Navier-Stokes equations shown in Equation 2. The second-order Crank-Nicolson scheme was employed for the diffusion and VPM terms. The other terms were calculated explicitly with the third-order Runge-Kutta scheme [40].

### Effect of the grid resolution on flow distribution

To verify the current grid resolution, additional numerical simulations with doubled grid resolutions in all the directions were performed. Due to the large computational cost, we considered one set of KO mice and their control littermates (CTLR) for each of the *Foxo1* and *Prkci* cases. They are denoted as *Foxo1* CTLR-1F, *Foxo1*^*i*ΔEC^-1F, *Prkci* CTLR-1F and *Prkci*^*i*ΔEC^-1F, respectively. The numbers of grid points employed for the simulations with finer grid size are listed in the bottom of Table1.

The distributions of the velocity intensity around the angiogenic front obtained with current and finer grid resolutions are compared in Fig. S1. The left column contains the results obtained with the initially coarse resolution, whereas the right column shows those with the finer resolution. In general, the overall trend does not change with increasing the grid resolution. Specifically, it can be confirmed that the flow rate decreases at the angiogenic front region of the *Foxo1*^*i*ΔEC^ models, whilst it is enhanced in the same region of the *Prkci*^*i*ΔEC^ structures in comparison to the corresponding control structures. More quantitatively, Table S1 summarizes the average flow portion (integrated flow rate in percentage) in the inner plexus and angiogenic front regions of the *Foxo1*^*i*ΔEC^ and *Prkci*^iΔEC^ cases and their controls, before and after grid refinement. In general, the simulation with the coarser grids tends to overestimate the averaged flow portion in the angiogenic region by around 3%. This could be explained by better prediction of flow distributions within small-diameter branching vessels in the inner plexus with the grid refinement. The decrease of the averaged flow rate in the angiogenic region due to *Foxo1* knockout is estimated as 9.2% for the coarse resolution, whereas 9.8% for the finer one. Similarly, knocking out *Prkci*^iΔEC^ increases the flow rate in the angiogenic region by 15.8% and 16.1% for the coarse and fine resolutions, respectively. Hence, it can be confirmed that further grid refinement from the initial coarse resolution has only minor impacts on the current results, and does not change the current conclusions.

### Statistical analyses

All statistical analyses were carried out using Prism software (Graph Pad, CA). A P<0.05 was considered statistically significant. Data are based on at least three independent experiments or three mutant and control animals for each stage.

## Acknowledgments

Funding for this project was provided by the German Research Foundation, Deutsche Forschungsgemeinschaft (GRK2213), and the JSPS KAKENHI Grant Number JP17H03170 and JP17KK0128

## Author contributions

Y.H., and M.N. designed the study. F.M., Y.K. and T. H., performed experiments. F.M., Y.K., Y.H., and M.N. wrote the manuscript.

## Data and materials availability

All data is available in the main text or the supplementary materials.

## Supplementary Materials

**Table S1.** The ratio of the averaged flow rates in the inner plexus and angiogenic front regions before and after grid refinement.

|  | Coarse resolution |  | Fine resolution |  |
| --- | --- | --- | --- | --- |
|  | Inner Plexus | Angiogenic front | Inner Plexus | Angiogenic front |
| *Foxo1* CTRLR | 75.0 % | 25.0 % | 77.8 % | 22.2 % |
| *Foxo1*<sup>iΔEC</sup> | 84.2 % | 15.8 % | 87.6 % | 12.4 % |
| *Prkci* CTRLR | 78.7 % | 21.3 % | 81.8 % | 18.2 % |
| *Prkci*<sup>iΔEC</sup> | 62.9 % | 37.1 % | 65.7 % | 34.3 % |

**Figure S1.**
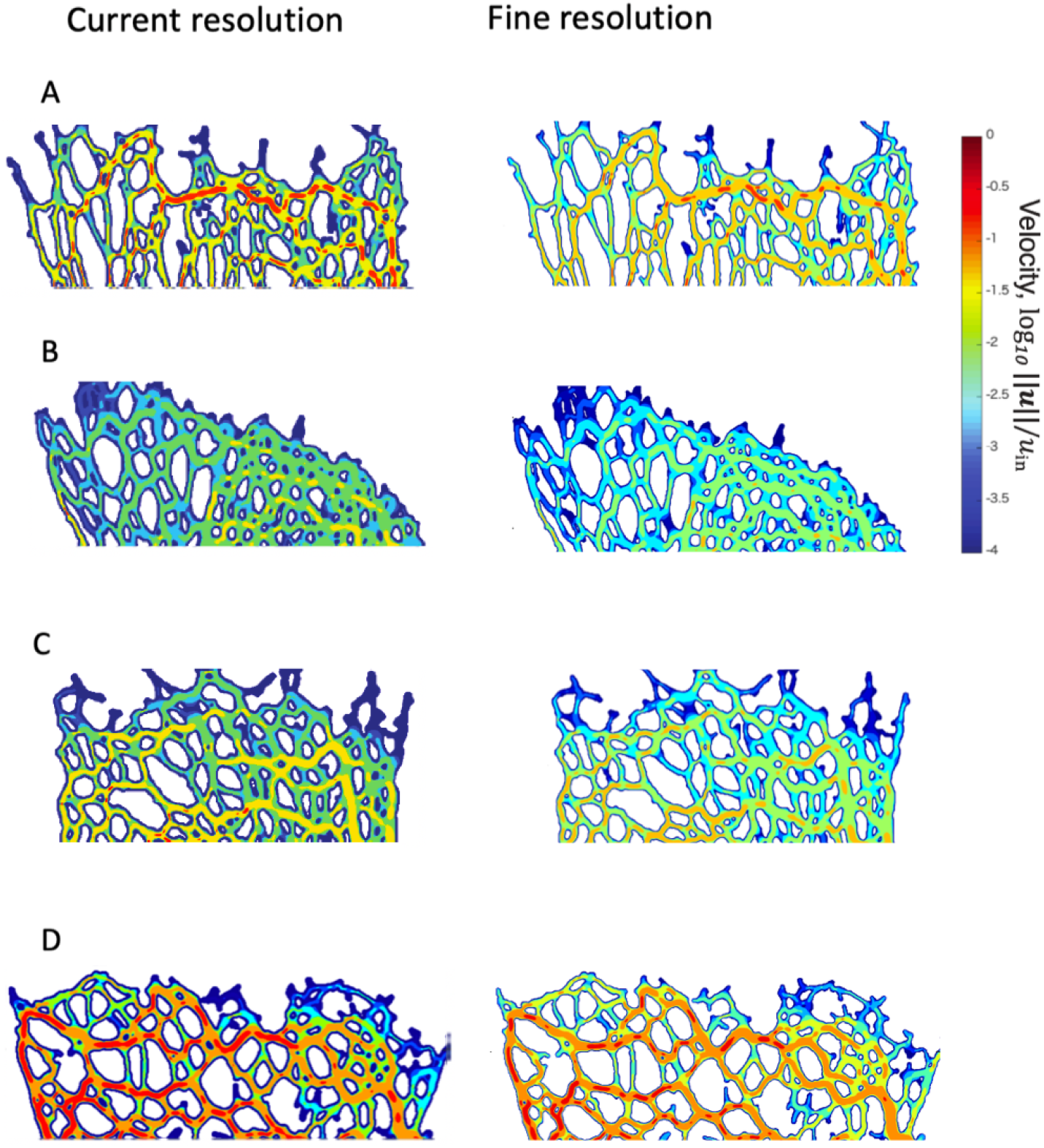
Visualization of the velocity distributions around the angiogenic front obtained with (left) current and (right) finer grid resolutions. A) *Foxo1* CTLR, B) *Foxo1*^iΔEC^, C) *Prkci* CTLR, D) *Prkci*^iΔEC^. In all figures, the local velocity intensity normalized by the inlet velocity is shown in a logarithmic scale.

